# Optimizing reaction coordinate by flux maximization in the transition path ensemble

**DOI:** 10.1101/2021.11.18.469181

**Authors:** Wenjin Li

## Abstract

Transition path ensemble is a collection of reactive trajectories, all of which largely keep going forward along the transition channel from the reactant state to the product one, and is believed to possess the information necessary for the identification of reaction coordinate. Previously, the full coordinates (both position and momentum) of the snapshots in the transition path ensemble were utilized to obtain the reaction coordinate (J. Chem. Phys. 2016, 144, 114103; J. Chem. Phys. 2018, 148, 084105). Here, with the conformational (or position) coordinates alone, it is demonstrated that the reaction coordinate can be optimized by maximizing the flux of a given coordinate in the transition path ensemble. In the application to alanine dipeptide in vacuum, dihderal angles *ϕ* and *θ* were identified to be the two best reaction coordinates, which was consistent with the results in existing studies. A linear combination of these two coordinates gave a better reaction coordinate, which is highly correlated with committor. Most importantly, the method obtained a linear combination of pairwise distances between heavy atoms, which was highly correlated with committor as well. The standard deviation of committor at the transition region defined by the optimized reaction coordinate is as small as 0.08. In addition, the effects of practical factors, such as the choice of transition path sub-ensembles and saving interval between frames in transition paths, on reaction coordinate optimization were also considered.

## Introduction

Molecular dynamics (MD) simulation is an molecular approach that is increasingly applied in the study of biomolecular systems at the atomic level and provides increased understanding of the underlying mechanisms.^1,2^ It seems a common consensus that the essential dynamics of these systems could be captured by a few coordinates, which are named as reaction coordinates. Based on such a physical viewpoint, many biological processes, such as protein folding, protein-ligand interactions, and conformational changes, can be simplified as rare transitions between two stable states that are well-defined by a reaction coordinate and are separated by high activation barriers.^3^ Most of the time the system stays in one of the stable states, while the transition from one stable basin to the other rarely occurs. As these biological processes resemble chemical reactions, the rare transitions are all reactive trajectories, which keep going forward to a great extent from one state (e.g., reactant state) to the other (e.g., product state). Usually, each reactive trajectory moves around and follows tightly the reaction coordinates, thus it is believed that a collection of all these reactive trajectories (or transition paths in the conformational space) possesses the information necessary for obtaining reaction coordinates, as well as the detailed mechanisms and kinetic information of the process.^4–7^ These reactive transitions or transition paths form a special ensemble named transition path ensemble (TPE). Since the time of reactive transitions is usually orders of magnitude faster than the time spent in the stable states, the computational cost in sampling of these transition paths is much cheaper than a brute-force sampling of an equilibrium trajectory. In the endeavour of sampling transition paths, transition path sampling (TPS),^4,8–11^ forward flux sampling, ^12,13^ nudged elastic-band method,^14,15^ and finite-temperature string method^16,17^ are representative path-finding approaches developed in the past decades. Among them, TPS proposed by Chandler and coworkers is worthy of particular attention. In TPS, transition paths are sampled without the priori knowledge of the reaction coordinate by Monte Carlo moves in the space of transition paths with an initially known transition path as a seed.^4,8–11^ The sampled paths are natural, as no bias potential is applied, and are properly weighted as in an equilibrium trajectory. TPS has been applied widely in studying complex biomolecular systems.^18–27^

With the TPE obtained from path-finding algorithms, extracting reaction coordinates out of the TPE is also challenging. Many early attempts to this challenge relies on additional information such as committor or equilibrium distribution of conformations. The committor of a given configuration is the probability of a trajectory initiated with Boltzmann distributed momenta to commit to the product state first^28–30^ and is widely regarded as the ideal reaction coordinate.^7^ Ma and Dinner^31^ proposed an automatic method that identifies reaction coordinates by utilizing the committor to train a machine learning model. Best and Hummer looked for a coordinate with sharpest peak in the distribution of a special probability, that is the probability of a trajectory to be a transition path provided that it visits configurations at a specific location of the coordinate, to be the reaction coordinate.^23^ In the method, one has to calculate the equilibrium probability distribution of configurations along the coordinate using enhanced sampling methods. Other committor-based methods includes likelihood maximization method and its extensions,^32–34^ transition state ensemble optimisation,^6,35^ and cross-entropy minimization method.^36^ Instead of TPE, several approaches for reaction coordinate identification that require either an equilibrium trajectory or an ensemble of non-equilibrium short trajectories were proposed as well. ^34,37,38^ With an ensemble of non-equilibrium short trajectories, the committor of configurations and thus the ideal reaction coordinate can be estimated by recently proposed algorithms such as Markov state models, ^39,40^ non-equilibrium non-parametric analysis,^41^ and iso-committor surfaces calculation.^42^ Machine learning methods were also frequently applied to find reaction coordinates for biomolecular systems without the information of committor.^43,44^

In the very recent endeavour to identify reaction coordinate from the TPE, the so-called emergent potential energy along a coordinate, a quantity estimated from the configurations of the TPE alone under the assist of forces on each atom (calculated by a rerun of the TPE without further sampling), was used to quantify the quality of a coordinate as the reaction coordinate and suggested a new energy decomposition scheme along the reaction coordinate.^45^ Later, the equipartition terms in the TPE were calculated from both the position and momentum coordinates of the reactive trajectories and were suggested to be a promising quantity to appraise reaction coordinate.^46^ In this paper, we are concerned with the extraction of reaction coordinate from the TPE with the conformational coordinates only. One attractive difference between TPE and equilibrium ensemble is that all the states in the TPE have the tendency to going towards the final state, while a state in the equilibrium ensemble has no tendency towards any state. Each transition path can be counted as a flux from the starting state to the final one, and the total or net flux in the TPE equals the number of transition paths. In the equilibrium ensemble, there will be equal number of trajectories that go out of and enter into a state, then the flux vanishes. One can imagine that the distribution of velocity for any coordinate in the equilibrium ensemble will be in perfect symmetry around zero, while there will be more positive velocities than the negative ones for the reaction coordinate in the TPE if we assume the value of the reactant state increases when going from the starting state to the final one. In another words, there is a positive flux along the reaction coordinate in the TPE and the fluxes along any coordinate in the equilibrium ensemble are zero. For coordinates that are not important for the transition (e.g., bath modes), the distributions of their velocities should resemble the equilibrium case and thus the fluxes along them are expected to be vanishingly small. Therefore, the flux along a coordinate in the TPE could be a promising quantity to measure the relevance of a coordinate to the reaction coordinate.

Here, we tested whether it is feasible to find and optimize reaction coordinates using the flux of a coordinate in the TPE in a well-studied system, that is the *C*_7eq_ → *C*_7ax_ isomerization of the alanine dipeptide in vacuum, of which the reaction coordinate is largely known.^31,36,45–48^ In the following, we start by introducing the formula of flux along a coordinate in the TPE. Next, the power of the flux as a quantity to appraise reaction coordinates is illustrated on the alanine dipeptide in vacuum. Then, a number of practical aspects is considered to show the promise of the proposed approach in its application to more complex systems. Finally, concluding remarks are given to end the paper.

## Method

### Flux along a coordinate

As all the transition paths start from the reactant state and largely keep going forward along the reaction coordinate to reach the product state, by counting the net number of transitions across a surface at a fixed location of the reaction coordinate, it will be equal to the number of total transition paths in the TPE. Here, we define that the total number of transitions (include both forward and backward transitions) across a surface at a specified position of a coordinate as the flux in a given ensemble. In case of the TPE, the flux at any value of the reaction coordinate equals the number of transition paths, while the flux at any position of a bath mode is vanishing small. As is shown in Fig. 1, two imaginary transition paths are projected on two coordinates, the reaction coordinate (RC) and a bath mode (q). Two stable basins (A and B) were defined by the reaction coordinate and the transition paths initiate from the boundary of state A (S_*A*_) and end at the boundary of state B (S_*B*_). The flux at a fixed location of the reaction coordinate (the vertical dotted line in red) is 2 as two transition paths cross the vertical line forwardly. On the other hand, the horizontal dotted line in red is a surface at a special position of the bath mode. The number of intersections in the forward direction for both transition paths equals the one in the backward direction and thus the flux cross the surface is zero. Thus, it is expected that the closer a coordinate is related to the reaction coordinate, the higher the flux along it is.

**Figure 1:**
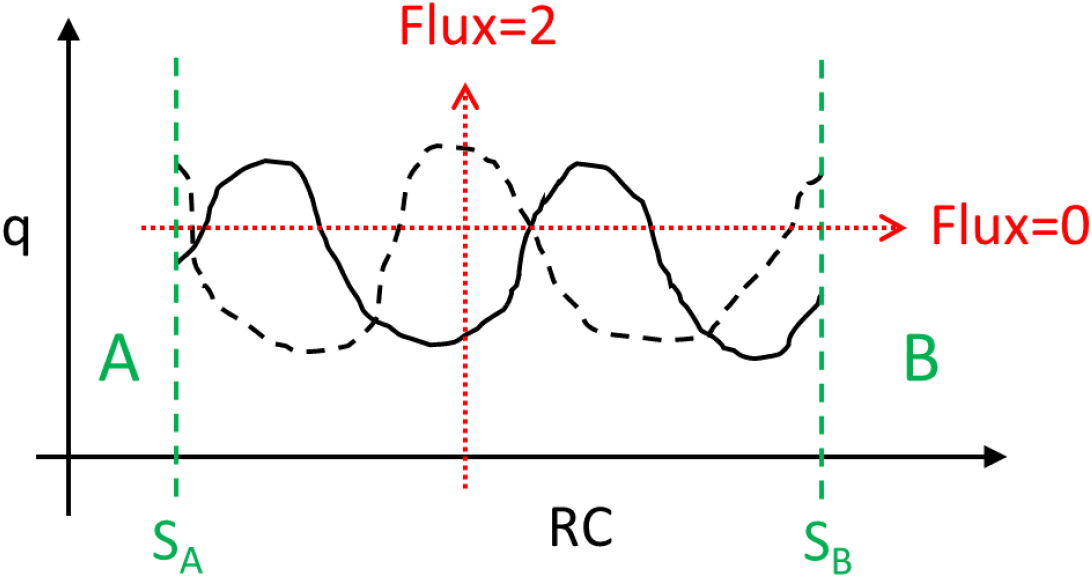
(schematic) Illustration of flux along two coordinates in the transition path ensemble. Two transition paths (solid and dashed lines in black) projected onto two coordinates, the reaction coordinate (RC) and a bath mode (*q*). S_*A*_ and S_*B*_ (green dashed lines) are the surfaces that defines the boundary of state A and state B, respectively. The flux at a specific location of both coordinates are indicated by red dotted lines.

For an ensemble of trajectories in a configuration space defined by ***q***, where ***q*** = (*q*_1_, *q*_2_, · · ·, *q*_n_) is the configurational coordinates of the system, the flux at the fixed location *ξ′* of a coordinate *ξ*(***q***), an arbitrary function of ***q***, is given by

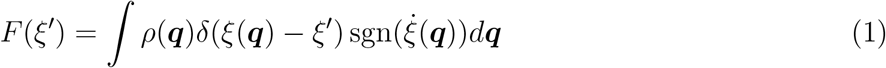

where, *ρ*(***q***) is the probability density of configuration ***q*** in the ensemble, 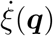 is the generalized velocity of *ξ*(***q***), *δ*(*x*) is the Dirac delta function, and sgn(*x*) is the sign function. Eq. 1 gives the flux as a function of a coordinate. To facilitate the comparison of fluxes between different coordinates, the flux along the coordinate is averaged with the probability distribution of configurations along the coordinate (*ρ*(*ξ*)) as weight. The averaged flux of a coordinate is thus defined as,

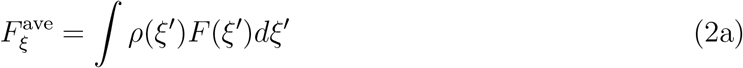

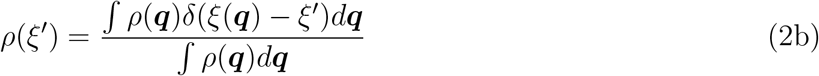

In practice, a trajectory from an MD simulation is usually saved as a string of frames with a fixed time interval and thus a time-series of *ξ*(*t*) as a function of time is obtained. From such a time series, the flux as a function of *ξ* is obtained by

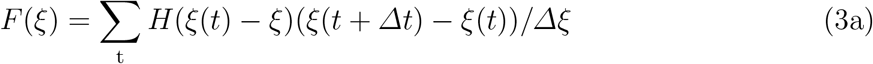

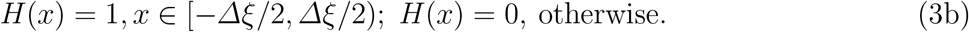

where, *Δξ* is the bin size for *ξ* in such an histogram-based approach, *Δt* is the trajectory saving interval. Analogously, the probability distribution of configurations along *ξ* is given by

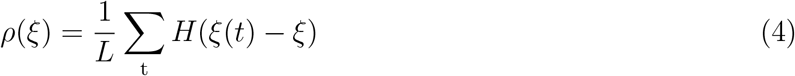

where, *L* is the total number of configurations in the ensemble of trajectories. Then the averaged flux along a coordinate is,

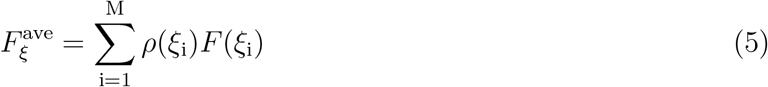

Here, *M* is the number of bins, and the summation is over all the bins in the histogram-based estimations of both the flux and the probability distribution along a coordinate. For the comparison of fluxes between different TPEs, the relative flux (RF) and the averaged relative flux (ARF) are used and are given by the flux and the averaged flux divided by the number of transition paths, respectively.

### Transition path ensemble

The TPE with 526 transition paths were harvested with the same simulation setup and definition of stable basins as detailed in Ref. [^48^]. The stable basins were defined based on dihedral angles *ϕ* and *ψ* (see Table S1 and Fig. S1 for their definition). Each transition path is 2 ps in length and contains 4000 configurations. The committors of every 10th configurations in a transition path were evaluated with 400 shooting trajectories. The committors of the rest configurations were derived with a linear interpolation.

The TPE were divided into several sub-ensembles, which consists of transition path segments as defined in Fig. 2. For instance, the transition path sub-ensemble S0 consists of segments S0. The transition barrier segment (TBS) was divided into 10 equal segments. The reactant segment (RS) and product segment (PS) were divided into 40 equal segments. The boundaries of state A and B (S_*A*_ and S_*B*_) can be defined to be the same as the one in transition path sampling, and the transition path sub-ensembles with such definitions were named with the prefix of “PHI”. S_*A*_ and S_*B*_ can also be defined based on the committor, and they are defined to be the surfaces at PB = 0.001 and PB = 0.999, respectively. The corresponding transition path sub-ensembles were named with the prefix of “PB”. For example, the transition path sub-ensemble S0 defined by the committor was named “PB0”

**Figure 2:**
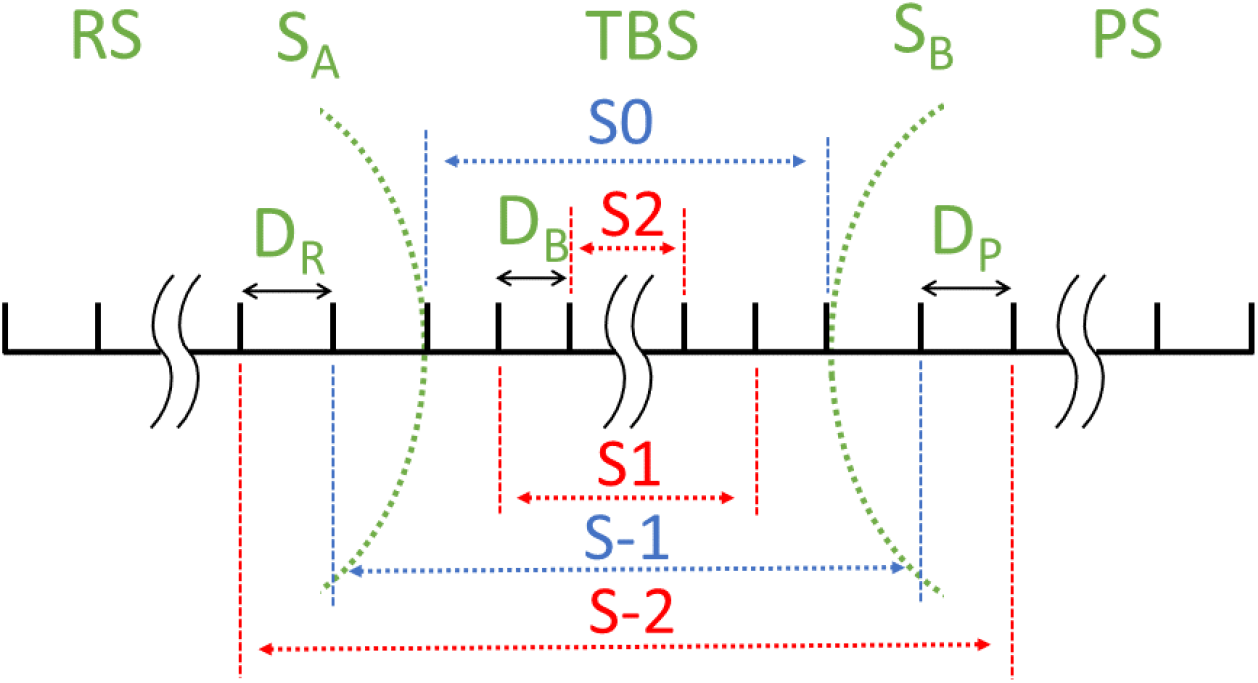
(schematic) The definition of segments in a transition path. The transition path is represented by a straight horizontal line in black, with vertical marks. S_*A*_ and S_*B*_ (green dashed lines) are the surfaces at the boundaries of states A and B, respectively. They divide the transition path into the reactant segment (RS), the transition barrier segment (TBS), and the product segment (PS). The TBS starts from where the transition path crosses last with S_*A*_ to where it crosses first with S_*B*_. RS and PS are the segments before and after TBS, respectively. These three segments are divided into several equal segments (D_*S*_, D_*B*_, and D_*P*_), as indicated by the vertical marks. The TBS is the segment S0; Removal of one segment D_*B*_ at both ends of the TBS gives the segment S1. Adding one segment D_*S*_ and one segment D_*P*_ before and after the TBS gives the segment S-1. Segments S2 and S-2 and others are defined in a similar way.

### Flux maximization as a linear combination of selected coordinates

For a given set of coordinates ***r*** = (*r*_1_, *r*_2_, · · ·, *r*_m_), which are functions of conformational coordinates ***q***, we aim to find a linear combination of these coordinates, of which the ARF is of the maximum magnitude. Let us assume that *ξ*° is a linear combination of ***r*** with coefficients 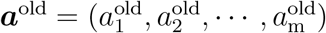, that is *ξ*^old^ = ***a***^old^ · ***r***, and *ξ*^new^ = *cosα* * *ξ*^old^ + *sinα* * *r*_i_, a linear combination of *ξ*^old^ and one of the coordinates in ***r***. By estimating the ARFs of *ξ*^new^ at different values of *α*, the value of *α* at which the ARF of *ξ*^new^ reaches its maximum in magnitude (*α*^max^) can be determined. In practice, ARFs at a finite set of points (e.g., 100 is used by default in this work) that are evenly distributed in the range of [0, *π*]. Then, *ξ*^new^ can be expressed as a linear combination of ***r*** with coefficients ***a***^new^, which is given by

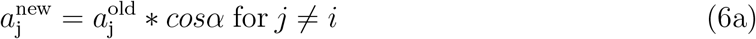

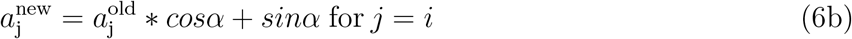

Starting with *ξ*^old^ being one of the coordinates in ***r***, for example the coordinate with the highest magnitude of ARF, one can select another coordinate in ***r*** as *r*_i_ and a new coordinate *ξ*^new^ can be obtained. *ξ*^new^ is then assigned to *ξ*^old^ and the above optimization process can be repeated with all the coordinate in ***r*** are used in turn. Such iterative process stops when the ARF of the optimized RC (*ξ*^new^) converges and results in an optimized RC as a linear combination of selected coordinates.

## Results and Discussions

### Flux in the TPE for BAT coordinates

We first considered a complete set of coordinates as a full description of the alanine dipeptide in vacuum. The set of coordinates consists of the 6 external coordinates and the so-called BAT internal coordinates (the bonds, angles and dihedral angles), the same as the ones used in previous studies^45,46,48^ (see Table S1). In such a coordinate set, one proper dihedral is used with the rest ones being improper dihedrals when there are two or more branches at an atom. The coordinate set possesses some invariant properties with respect to the choice of the BKS-tree.^49,50^ In the MD simulation of the *C*_7eq_ → *C*_7ax_ isomerization of the alanine dipeptide in vacuum, the translational and rotational motions of the system were removed. Thus, these 6 external coordinates are believed to be trivial and only the 60 internal coordinates are considered in following analyses. As the committor is the ideal reaction coordinate, the sub-ensemble S0 (as defined in Fig. 2) of the transition paths defined by the committor are used as the standard sub-ensemble for testing the performance of the flux as a quantity to appraise reaction coordinates (Later, we will show that the committor information is not required for obtaining the reaction coordinate). The ARFs along the 60 internal coordinates are shown in Fig. 3A. For most of the coordinates, their ARFs are close to zero. As ranked by the magnitude of ARFs, the top four coordinates are *ϕ*, *θ*, the 48th coordinate (*dih*_7_6_4_8), and *ψ*, as listed in a decreasing magnitude of ARFs (see Table 1). Thus, the *ϕ* and *θ* are suggested to be the most importance coordinates, a result that is consistent with several existing studies. ^31,36,45–48^ The ARFs of *ϕ* and the 48th coordinate are positive, while the ones of *θ* and *ψ* are negative. The coordinates with non-zero ARFs are mainly dihedral coordinates and the ARFs of almost all the bonds and angles are largely zero. The RFs along each coordinate were also examined. It is not surprising to observe that the RFs of all the bonds and angles (except the 5th and 35th coordinates) at any location of the coordinate are close to zero (see Fig. S2). For several dihedral coordinates with non-zero ARFs, the magnitudes of RFs along them first increase from zero and then decrease to be close to zero, while the RFs of most dihedral coordinates are largely zeroes along the coordinate (see Fig. 3B). An exception is the flux along the 54th coordinate, whose flux is non-zero and of almost the same magnitude at any location. Such an abnormal observation could be due to the insufficient sampling along this coordinate. Nevertheless, it is safe to conclude that the 54th coordinate is a trivial coordinate.

**Figure 3:**
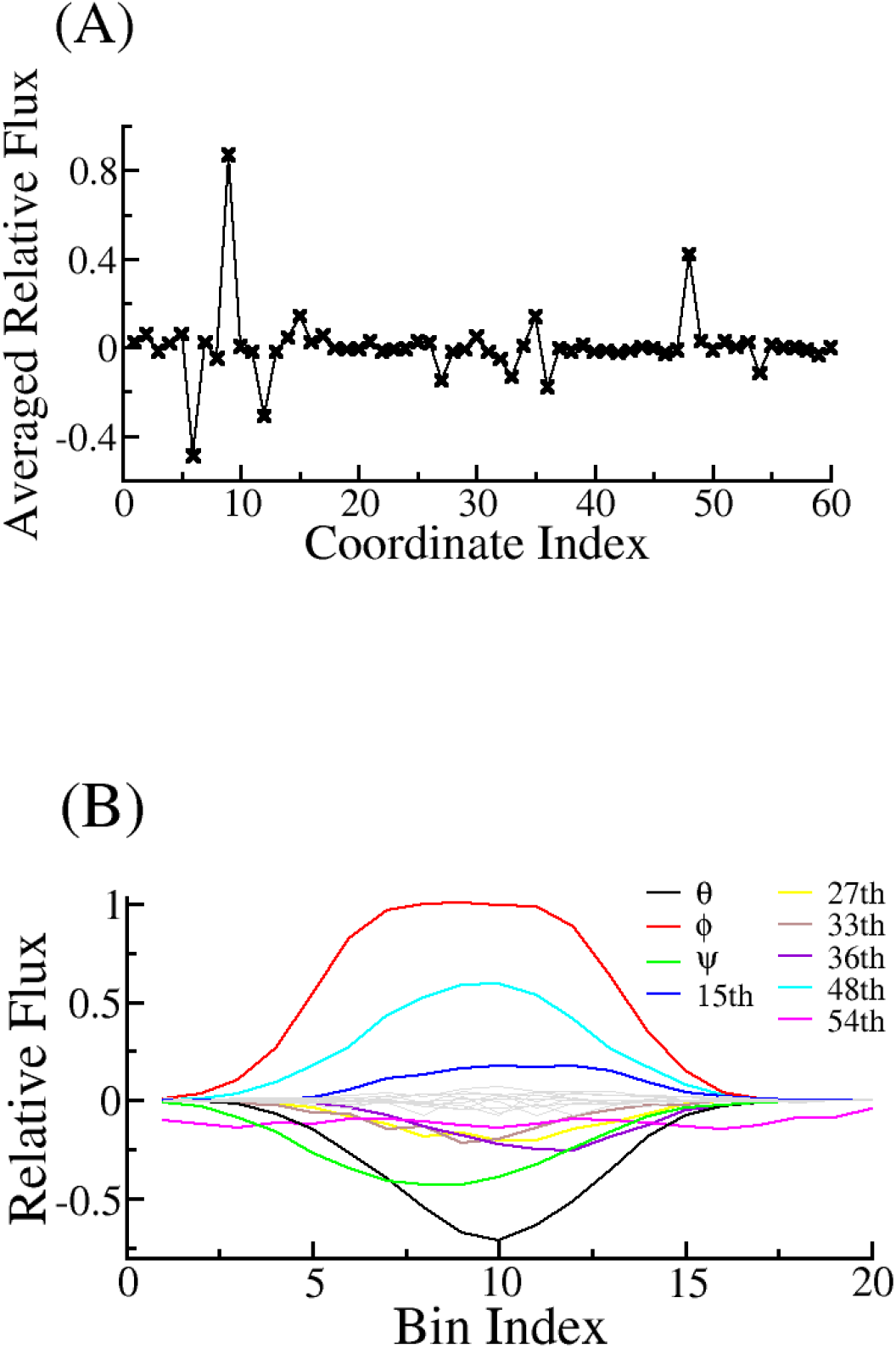
Relative fluxes of internal BAT coordinates in the transition path ensemble defined by the committor. (A) The averaged relative fluxes for the 60 internal coordinates (*M* = 400). (B) The relative fluxes along the dihedral coordinates (*M* = 20). Single peak or plateau appears in the fluxes along several coordinates, such as *ϕ*, *θ*, and *ψ*. The relative fluxes along other coordinates are close to zero and are shown in grey lines. The flux of the 54th coordinate is likely to be not properly estimated. See Table S1 for the indexes of the coordinates.

**Table 1:** Information of important internal BAT coordinates. Averaged relative flux (ARF), the rank by the magnitude of ARF (rank1), the coefficient in the optimized reaction coordinate (coef.), and the rank by the absolute value of the coefficient (rank2) for each coordinate are listed. Coordinates in the top 8 of either rank1 or rank2 are listed. The full list and detailed description can be found in Table S1.

| index | name | ARF | rank1 | coef. | rank2 |
| --- | --- | --- | --- | --- | --- |
| 5 | ang_C4_N6_CA8 | 0.06 | 11 | -0.032 | 8 |
| 6 | dih_O5_C4_N6_CA8 ($\theta$) | -0.49 | 2 | 0.477 | 2 |
| 9 | dih_C4_N6_CA8_C14 ($\phi$) | 0.869 | 1 | 1 | 1 |
| 12 | dih_N6_CA8_C14_N16 ($\psi$) | -0.312 | 4 | 0.044 | 6 |
| 15 | dih_CA8_C14_N16_CH18 | 0.14 | 7 | -0.062 | 5 |
| 27 | dih_H17_N16_C14_CH18 | -0.149 | 6 | -0.027 | 9 |
| 35 | ang_N6_CA8_CB10 | 0.138 | 8 | -0.037 | 7 |
| 36 | dih_CB10_CA8_N6_C14 | -0.179 | 5 | 0.064 | 4 |
| 48 | dih_H7_N6_C4_CA8 | 0.421 | 3 | -0.024 | 10 |
| 51 | dih_CH1_C4_O5_N6 | 0.024 | 24 | -0.084 | 3 |

Interestingly, the flux along *ϕ* presents a plateau, where the RF is 1 and thus the flux equals the number of transition paths. One would expect that the RF along a reaction coordinate will be 1 everywhere, and thus gives ARF of 1. Therefore, the next step is to find a coordinate that is better than *ϕ* as the reaction coordinate. From the above analysis, it seems reasonable to hypothesize that the higher the ARF is, the better the coordinate as a reaction coordinate. Thus, we can optimize the reaction coordinate by maximizing its ARF. As a first attempt, we estimated the ARFs of coordinates as a linear combination of *ϕ* and *θ*, that is *RC* = *cosα* * *ϕ* + *sinα* * *θ* with different values of the angle *α*, and the obtained ARF as a function of *α* is plotted in Fig. 4. The maximum of the ARF is reached at *α*=0.19*π*, where the ARF is 0.916 (the ARF of *ϕ* is 0.869) and the optimized reaction coordinate is *RC*^opt^ = 0.83*ϕ* + 0.56*θ*, which is in close consistent with the ones proposed in existing studies (0.78*ϕ* + 0.63*θ* in ref. [^45^] and 0.81*ϕ* + 0.59*θ* in ref. [^46^]). As shown in Fig. 5A, the committor of configurations at any position of the optimized reaction coordinate is narrowly distributed and the averaged committor is a good sigmoid function of the reaction coordinate. The committors of configurations at *RC*^opt^ ∈ [0, 0.02] are narrowly distributed around 0.5 with standard deviation of 0.13 (Fig. 5B). As a direct comparison, the standard deviation of committors of configurations around the transition state defined by *ϕ* is 0.20 (see Fig. S3).

**Figure 4:**
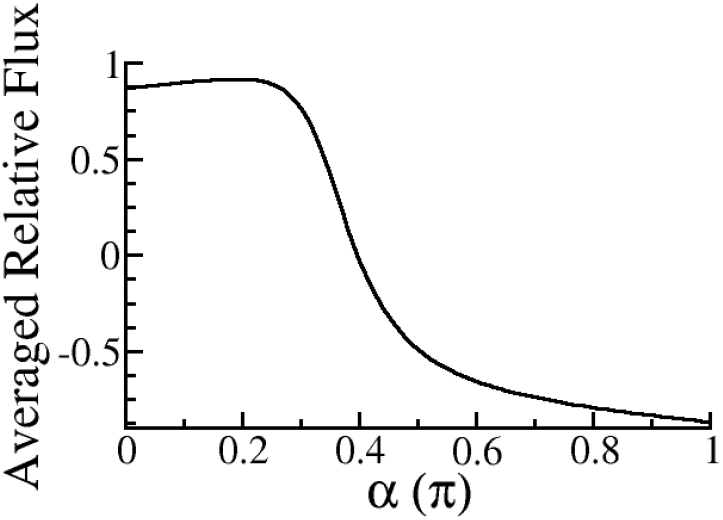
The averaged relative fluxes of the coordinate *RC* = *cosα ϕ* + *sinα θ* at different values of *α*.

**Figure 5:**
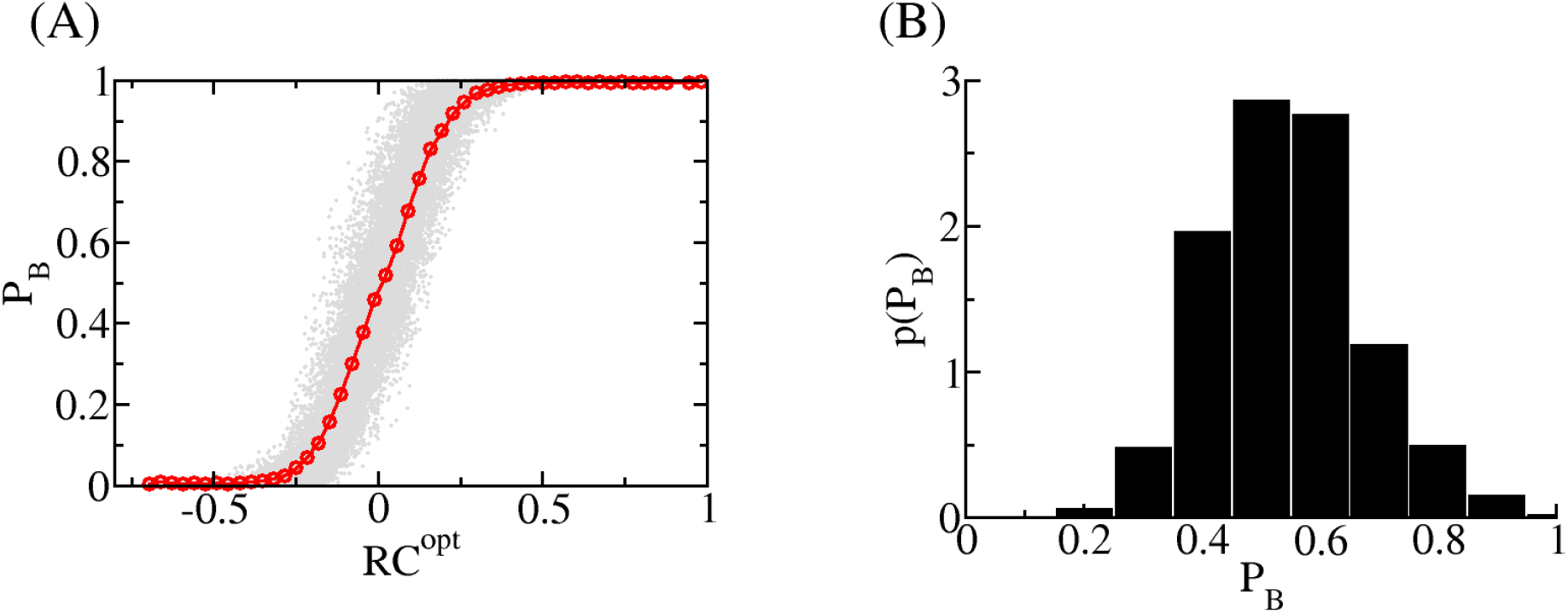
Committor analyses of an optimized reaction coordinate *RC*^opt^ = 0.83*ϕ* + 0.56*θ* (in the unit of radian). (A) The averaged committor of the configurations at fixed location of *RC*^opt^ as a function of *RC*^opt^ (red line with circles). The committors of all configurations are shown as circles in grey. (B) Histogram of the committors for configurations at *RC*^opt^ ∈ [0, 0.02].

Then we tried to optimize the reaction coordinate as a linear combination of all the 60 internal coordinates with the iteration method as detailed in the method part. The optimized flux is 0.932 and the committor of configurations is monotonically increasing with the optimized RC (Fig. 6A), with the distribution of *P*_B_ peaked at 0.5 and a standard deviation of 0.11 at the transition regions (Fig. 6B). In order to compare the contributions of each coordinate to the optimized RC, all the coordinates were normalized by their maximum and minimum values in the sub-ensemble of TPE and thus they are between 0 and 1. The magnitudes of coefficients of the coordinates in the linear combination are thus to some extent reflect their importances to the optimized RC. As can be seen from Table 1, dihedral angles *ϕ* and *θ* are ranked in the top 2 based on both ARF and coefficient in the optimized RC. For most of the coordinates with high magnitudes of ARFs, they are also ranked to be important by their coefficients in the optimized RC. Although the 48th coordinate is ranked No. 3 according to its ARF, it is ranked No. 10 based on its coefficient. However, by optimizing RC as a linear combination of *ψ* and the 48th coordinate, it gave *RC*^opt^ = 0.88*ϕ* − 0.48 * 48*th*, in which the coefficient of the 48th coordinate is comparable to the one of *θ* when the optimized RC is a linear combination of *ϕ* and *θ*. Since atoms to define the 48th coordinate and *θ* are on the same peptide plane, they are likely to be highly correlated and the contribution of the 48th coordinate to the optimized RC is largely taken into account by the including of *θ*, which results in a small coefficient for the 48th coordinate. On the other hand, the 51th coordinate on the same peptide plane with *θ* and the 48th coordinate is ranked No. 51 according to its ARF, which is as small as 0.024, it is ranked No. 3 based on its coefficient, which is though about one order of magnitude smaller the ones of the top 2 coordinates.

**Figure 6:**
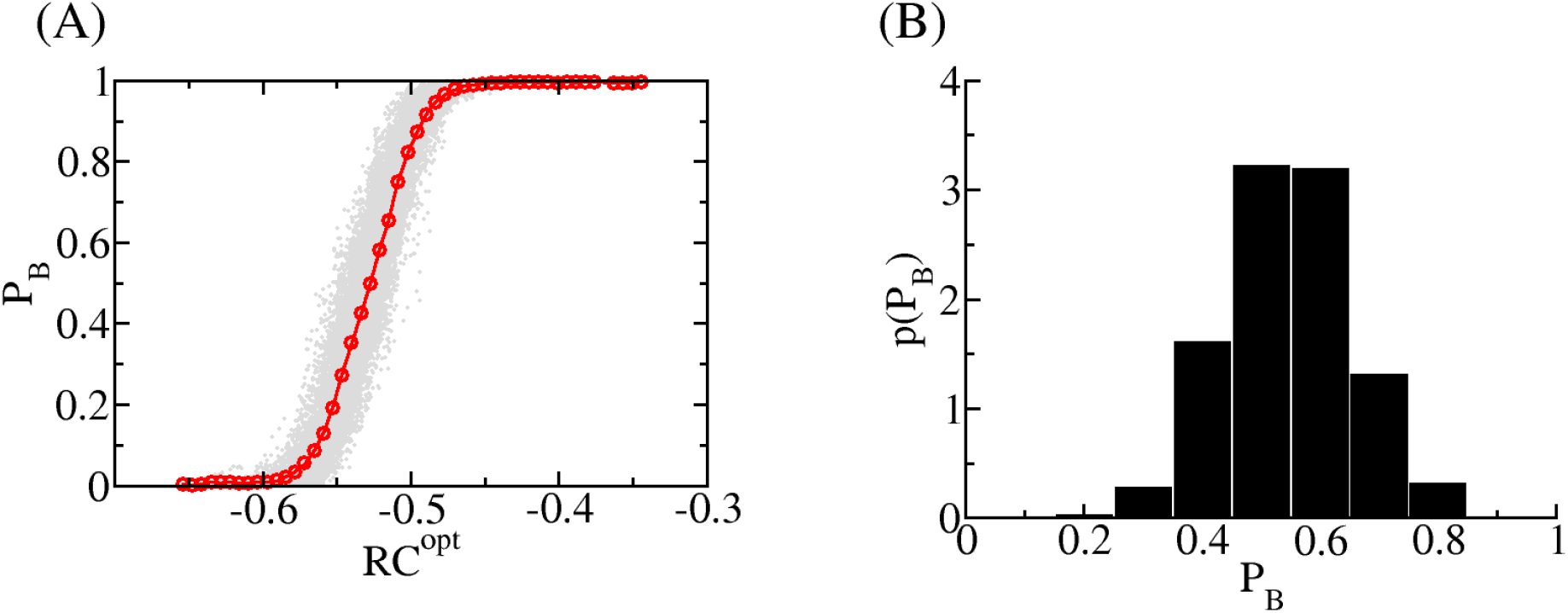
Committor analyses of an optimized reaction coordinate as a linear combination of internal BAT coordinates. (A) The averaged committor of the configurations as a function of the optimized reaction coordinate (red line with circles). The committors of all configurations are shown as circles in grey. (B) Histogram of the committors for configurations with the optimized RC between −0.532 and −0.524.

### Optimized RC as a linear combination of pairwise distances

For the study of biomolecules, such as protein and DNA, distances between atoms or residues are commonly used to construct the reaction coordinate.^51,52^ We thus tested whether a good reaction coordinate could be obtained from a set of coordinates that consists of the 45 pairwise coordinates between heavy atoms of the alanine dipeptide (see Table S2 for the list of pairwise coordinates). The ARFs along these coordinates are shown in Fig. 7. The coordinate with maximum ARF in magnitude is the distance between atoms O5 and CB10 (the 20th coordinate). It makes sense as the transition from *C*_7eq_ to *C*_7ax_ is largely hindered by the steric clash between atoms O5 and CB10. The averaged committor along the *O*5 − *CB*10 distance is plotted in Fig. S4 and is observed to be monotonically decreasing to a great extent, indicating that the *O*5 − *CB*10 distance is of satisfying quality as a reaction coordinate. The committors of configurations around the transition state defined by the *O*5 − *CB*10 distance is rather widely peaked at 0.5 with a standard deviation of 0.23. Thus, none of the pairwise distances is better than *ϕ* to be a reaction coordinate.

**Figure 7:**
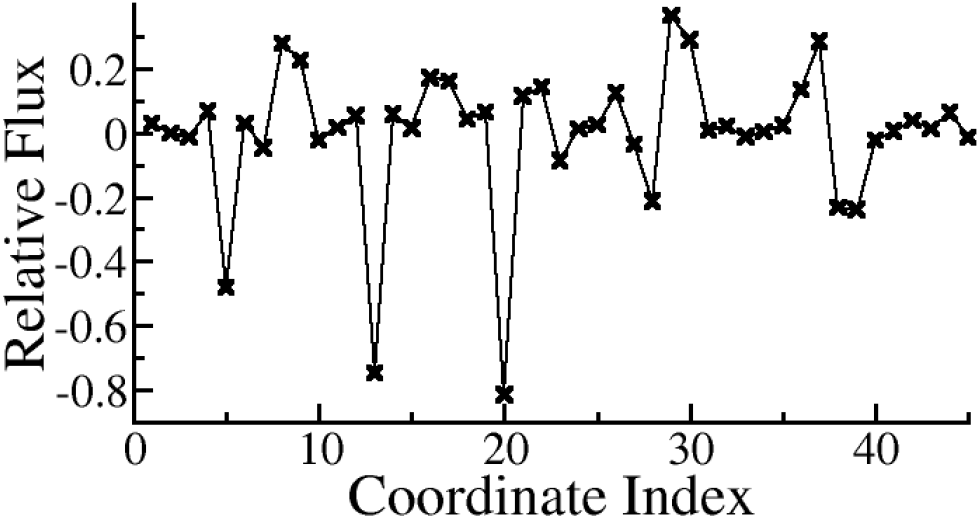
The averaged relative fluxes for the 45 pairwise coordinates in the transition path ensemble defined by the committor (*M* = 100). See Table S2 for the indexes of the coordinates.

Then we tried to optimize the reaction coordinate as a linear combination of these pairwise coordinates. The optimized flux is 0.928 and the committor of configurations is again observed to be monotonically increasing with the optimized RC (Fig. 8A), with the distribution of *P*_B_ narrowly peaked at 0.5 with a standard deviation of 0.08 at the transition regions (Fig. 8B). In analogue to the case of the internal BAT coordinates, all the coordinates were rescaled based on their maximum and minimum values in the sub-ensemble of TPE. As can be seen from Table 2, only half of the coordinates in the top 8 as ranked by their ARFs are also ranked in the top 8 according to their coefficients in the optimized RC. It could be due to the great redundancy in the set of pairwise distances. For example, all the top 3 coordinates (*O*5 − *CB*10, *C*4 − *CB*10, and *CH*1 − *CB*10) based on ARF describe to some extent describe the rotation of a peptide plane (similar to *ψ*) and thus contain redundant information. It also explains why the No. 2 and No. 3 coordinates by ARF are ranked No. 7 and 6 by coefficient, respectively. Two low ranking coordinates (*O*5 − *CA*8 and *C*4 − *C*14) by ARFs are of high coefficients, indicating that they contain information independent to the one in *O*5 − *CB*10.

**Figure 8:**
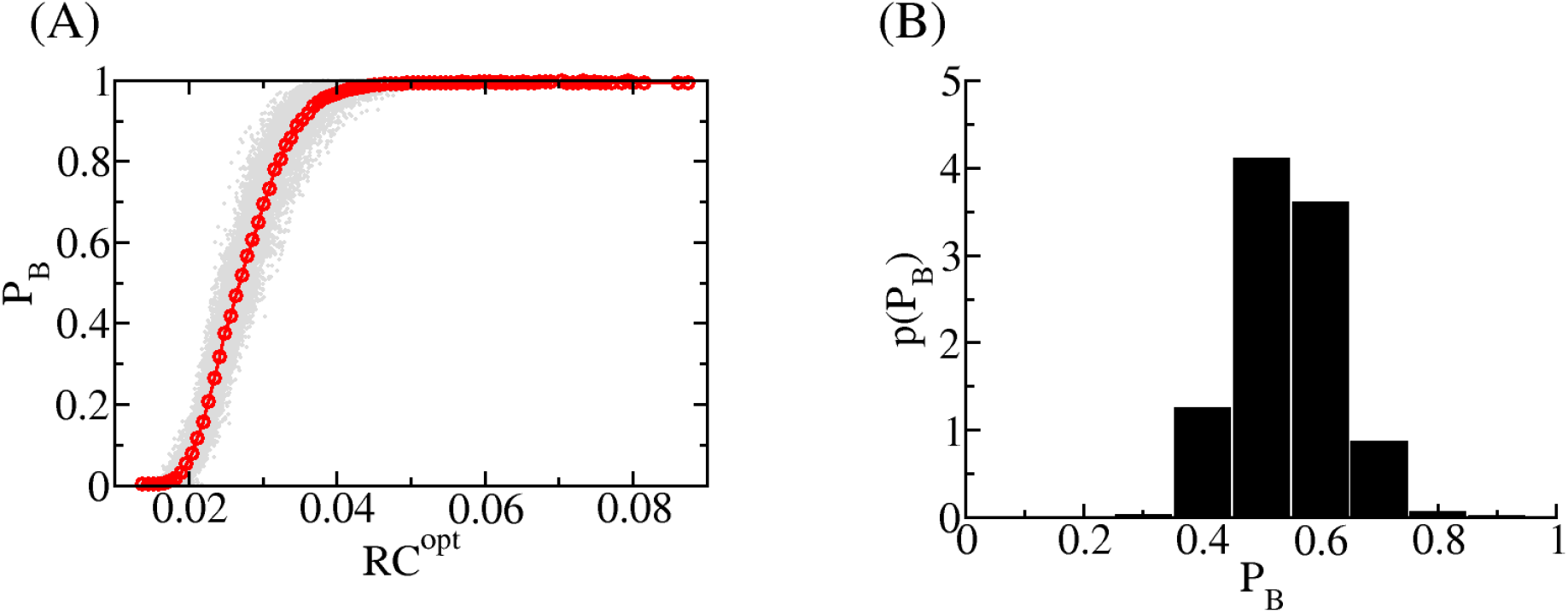
Committor analyses of an optimized reaction coordinate as a linear combination of pairwise distances. (A) The averaged committor of the configurations as a function of the optimized reaction coordinate (red line with circles). The committors of all configurations are shown as circles in grey. (B) Histogram of the committors for configurations with the optimized RC between 0.0264 and 0.0271.

**Table 2:** Information of important pairwise distances. Averaged relative flux (ARF), the rank by the magnitude of ARF (rank1), the coefficient in the optimized reaction coordinate (coef.), and the rank by the absolute value of the coefficient (rank2) for each coordinate are listed. Coordinates in the top 8 of either rank1 or rank2 are listed. The full list and detailed description can be found in Table S2.

| index | name | ARF | rank1 | coef. | rank2 |
| --- | --- | --- | --- | --- | --- |
| 5 | CH1-CB10 | -0.482 | 3 | -0.098 | 6 |
| 8 | CH1-N16 | 0.278 | 7 | -0.119 | 4 |
| 13 | C4-CB10 | -0.748 | 2 | -0.097 | 7 |
| 14 | C4-C14 | 0.058 | 22 | 0.144 | 3 |
| 19 | O5-CA8 | 0.064 | 20 | -0.184 | 2 |
| 20 | O5-CB10 | -0.815 | 1 | 1 | 1 |
| 26 | N6-CB10 | 0.123 | 16 | -0.112 | 5 |
| 29 | N6-N16 | 0.366 | 4 | 0.042 | 17 |
| 30 | N6-C18 | 0.289 | 5 | -0.042 | 15 |
| 36 | CB10-C14 | 0.134 | 15 | -0.085 | 8 |
| 37 | CB10-O15 | 0.285 | 6 | 0.066 | 12 |
| 39 | CB10-C18 | -0.24 | 8 | -0.072 | 10 |

### Several practical aspects

Since the ARF is estimated via a histogram-based method (Eq. 5), one concern is that whether the calculated ARF is sensitive to the number of bin (*M*). We thus obtained the ARFs of *ϕ* with different *M* and the result is shown in Fig. S5A. Although the calculated ARF of *ϕ* changes as *M* increases, it reaches a plateau and is largely independent to *M* when *M* is large enough (e.g., *M >* 10). The estimate ARFs of the internal BAT coordinates are largely consistent at M=50,100, and 400 (see Fig. S5B), thus we believe that the obtained ARFs are independent to the number of bin used in Eq. 5.

We further tested the dependence of the results on the choice of transition path sub-ensembles. The ARFs of internal BAT coordinates for sub-ensembles PB-1, PB0, and PB1 were calculated and shown in Fig. S6A. The ARFs of these sub-ensembles rank the coordinates in almost the same order. Although the absolute values of ARFs for coordinates *ϕ*, *ψ*, and *θ* are getting smaller as the transition path in the sub-ensemble become shorter (for the magnitudes of ARFs of these coordinates, PB-1*>*PB0*>*PB1), the optimized RCs as a linear combination *ϕ* and *θ* are considered to be almost identical (see Table 3) for these sub-ensembles.

**Table 3:** The ARFs and optimized RC obtained in different sub-ensembles. (see Fig. 2 for their definitions)

| sub-ensemble | ARF of $\phi$ | ARF of $\theta$ | ARF of 48th | ARF of $\psi$ | Optimized RC |
| --- | --- | --- | --- | --- | --- |
| PB-1 | 0.893 | -0.523 | 0.457 | -0.415 | $0.82\phi + 0.57\theta$ |
| PB0 | 0.869 | -0.490 | 0.421 | -0.312 | $0.83\phi + 0.56\theta$ |
| PB1 | 0.796 | -0.410 | 0.342 | -0.216 | $0.84\phi + 0.54\theta$ |
| PHI0 | 0.953 | -0.581 | 0.493 | -0.431 | $1.00\phi - 0.03\theta$ |
| PHI1 | 0.858 | -0.463 | 0.428 | -0.297 | $0.91\phi + 0.42\theta$ |
| PHI2 | 0.759 | -0.374 | 0.271 | -0.194 | $0.88\phi + 0.48\theta$ |

Since sub-ensembles discussed so far are defined based on the information of committor, which is quite computationally expensive to obtain, we thus test the proposed method on sub-ensembles defined based on the same stable basins as the one used in transition path sampling, that is based on the dihedral angles *ϕ* and *ψ*. The ARFs of internal BAT coordinates for sub-ensembles PHI-1, PHI0, PHI1, and PHI2 were calculated and shown in Fig. S6B. The ARFs of these sub-ensembles rank the coordinates in a largely consistent way, from which the same set of important coordinates can be identified. As noticed in a previous study,^48^ the behavior of *θ* in the transition region is significantly different from the one in the stable basins, here we observed that the magnitude of ARFs of *θ* in sub-ensemble PHI-1 is smaller than the one in sub-ensembles PHI0, while it increases from PHI2, PHI1, to PH0. The magnitude of ARFs of *θ* is comparable to the one of *ψ*. Thus, *θ* is a good reaction coordinate only in the transition region, a phenomenon observed previously as well.^48^ We thus focus on sub-ensembles PHI0, PHI1, and PHI2, and tested whether a good reaction coordinate can be obtained from such sub-ensembles. The optimized RCs as a linear combination *ϕ* and *θ* for sub-ensembles PHI0, PHI1, and PHI2 are shown in Table 3, which are consistent with the ones identified with sub-ensembles based on the committor with the exception for sub-ensemble PHI0. Since the sub-ensemble PHI0 is defined directly based on *ϕ* and *ψ*, it gives an ARF of *ϕ* as high as 0.953, which should be considered as an unusual results. Fortunately, the results from PHI1 and PHI2 are consistent very well with the ones when the committor is used to define the sub-ensembles. The obtained reaction coordinate can be used to define the stable basins and thus new sub-ensembles, from which the quality of the reaction coordinate can be further improved. Thus, a good reaction coordinate can be obtained from the TPE in a committor-free way.

For the alanine dipeptide in vacuum, the system consists of just 22 atoms and the transition occurs within 2 ps, it is thus possible to save every frames in the transition path. However, for complex biomolecular systems, it is not practical to save all the configurations in a transition path. We thus wonder whether the method can be applied to the cases when the transition paths are saved less frequently. For this purpose, the ARFs in the sub-ensemble PB0 were calculated when the transition paths were saved with time interval *Δ*t=0.5, 5, and 40 fs (see Table 4), which corresponding to the case of writing the trajectory file every, every 10th, and every 80th frame, respectively. For *Δ*t=0.5, 5, and 40 fs, the ARFs and their orders of top four coordinates, that is *ϕ*, *θ*, 48th coordinate, and *ψ* are largely consistent. Most importantly the optimized RC as a linear combination of *ϕ* and *θ* are almost the same(see Table 4). When *Δ*t=40 fs, there are 50 frames in each transition path and there are on average 5.4 frames per transition path in the transition barrier region. Thus, the approach is believed to be applicable to the TPE of complex biomolecules, in which the transition paths are usually saved infrequently.

**Table 4:** The ARFs and optimized RC when the TPE (PB0) is saved with different time intervals.

| $\Delta t$ (fs) | ARF of $\phi$ | ARF of $\theta$ | ARF of 48th | ARF of $\psi$ | Optimized RC |
| --- | --- | --- | --- | --- | --- |
| 0.5 | 0.869 | -0.490 | 0.421 | -0.312 | $0.83\phi + 0.56\theta$ |
| 5 | 0.879 | -0.495 | 0.435 | -0.313 | $0.86\phi + 0.51\theta$ |
| 40 | 0.929 | -0.510 | 0.435 | -0.321 | $0.80\phi + 0.59\theta$ |

## Concluding Remarks

Every transition path in a TPE can be viewed as a flux from the reactant state to the product one. The fluxes along coordinates that are relevant to the transition are thus believed to be non-zero and the reaction coordinate is assumed to be the one along which the flux is maximum. Here, we have described a committor-free approach to identify and construct the reaction coordinate by optimizing the flux along a proposed coordinate in the TPE. The method utilizes the conformational coordinates of the TPE alone and avoids the estimation of committor, a computationally expensive step. In its application to a well-studied biomolecular system, the *C*_7eq_ → *C*_7ax_ isomerization of the alanine dipeptide in vacuum, the same two coordinates, that is the dihedral angles *ϕ* and *θ*, as the ones reported in existing studies^45,46^ were identified to be the two most important coordinates. Importantly, the approach can construct an optimized RC as a linear combination of a coordinate set. Two different set of coordinates were examined: the internal BAT coordinates and the pairwise distances between heavy atoms. For both coordinate sets, a good reaction coordinate was obtained as a linear combination of all the coordinates in each set. The committor along the optimized RC is narrowly distributed (with standard deviation as small as 0.08 for the case of pairwise distances) and increases monotonically. Although the results depend on the definition of stable basins, it is demonstrated to be able to eliminate such dependence by dividing the TPE into different sub-ensembles. An iterative procedure can also be applied when necessary. It is also demonstrated that the approach can be applied to the case when the transition paths were saved with large time intervals. We thus believe that the approach is quite robust and is applicable to complex biomolecular systems.

## Supplementary Material

See supplementary material for the supplemental Figures S1-S6 and Tables S1-S2.

## Acknowledgments

This work was supported by Natural Science Foundation of Guangdong Province, China (Grant No. 2020A1515010984) and the Start-up Grant for Young Scientists (860-000002110384), Shenzhen University.

