## supplemental Figures and Tables for "Optimizing reaction coordinate by flux maximization in the transition path ensemble"

Wenjin Li
Institute for Advanced Study, Shenzhen University, Shenzhen, China

### Supplementary Material

#### Contents:

- Supplemental Figures and Tables: Figures S1-S6, Tables S1-S2.

### Supplemental Figures

S1 Schematic representation of the alanine dipeptide . . . . . . . . . . . . . . . . . . . . . S2
S2 The fluxes along the bond and angle coordinates . . . . . . . . . . . . . . . . . . . . . . . S2
S3 Committor analyses of $\phi$ . . . . . . . . . . . . . . . . . . . . . . . . . . . . . . . . . . . . S3
S4 Committor analyses of the $O5 - CB10$ distance . . . . . . . . . . . . . . . . . . . . . . S3
S5 The averaged relative flux estimated with different number of bins . . . . . . . . . . . S4
S6 The averaged relative flux of internal BAT coordinates for different transition path sub-ensembles . . . . . . . . . . . . . . . . . . . . . . . . . . . . . . . . . . . . . . . . . . . . S4

### Supplemental Tables

S1 Information of all the internal BAT coordinates . . . . . . . . . . . . . . . . . . . . . . S5
S1 Table S1 continued . . . . . . . . . . . . . . . . . . . . . . . . . . . . . . . . . . . . . . . . S6
S2 Information of all the pairwise distances . . . . . . . . . . . . . . . . . . . . . . . . . . . S7

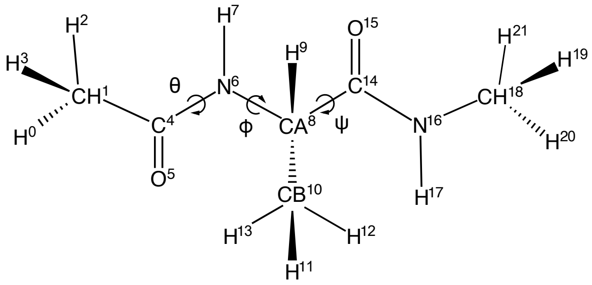


Figure S1: Schematic representation of the alanine dipeptide with indicated atom types and indices. The atoms are indexed from 0 to 21 and the indices are shown as the superscripts of the corresponding atom types. The atom is thus named as atom type followed by its index. For example, atom "CA8" is the atom of atom type CA and index 8. Three backbone dihedrals $\phi$, $\psi$, and $\theta$ are labelled (For their definitions, see Table S1).

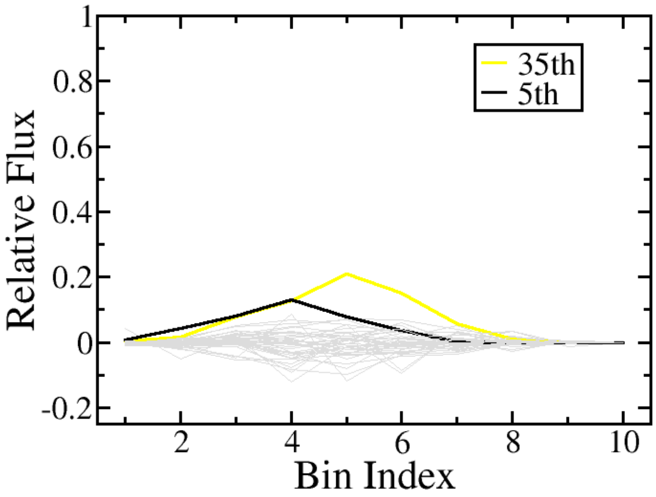


Figure S2: The fluxes along the bond and angle coordinates ($M = 10$). Single peaks appear in the fluxes along the 35th and 5th coordinates. The relative fluxes along other coordinates can be considered to be close to zero and are shown in grey.

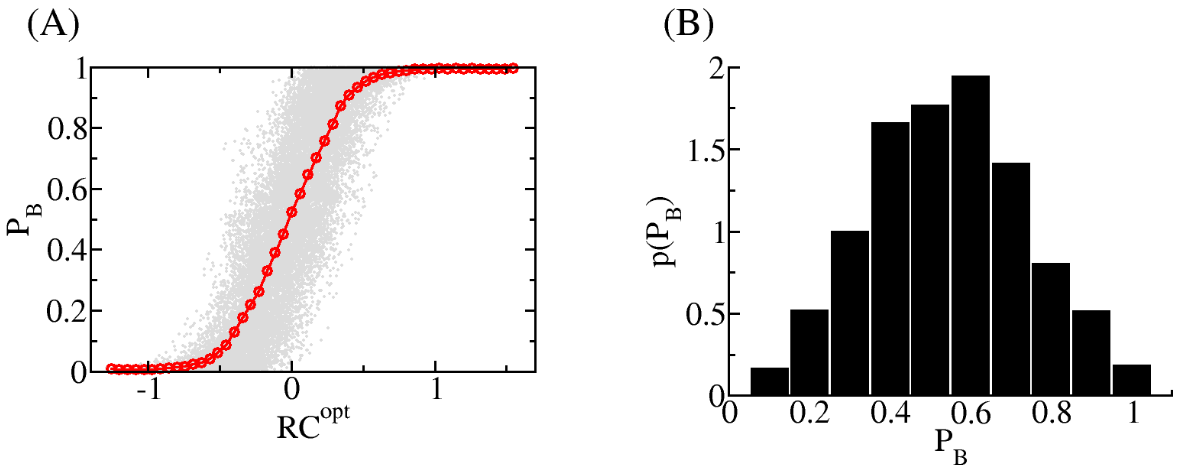


Figure S3: Committor analyses of $\phi$ (in the unit of radian). (A) The averaged committor of the configurations at fixed location of $\phi$ as a function of $\phi$ (red line with circles). The committors of all configurations are shown as circles in grey. (B) Histogram of the committors for configurations at $\phi \in [-0.06, 0.02]$.

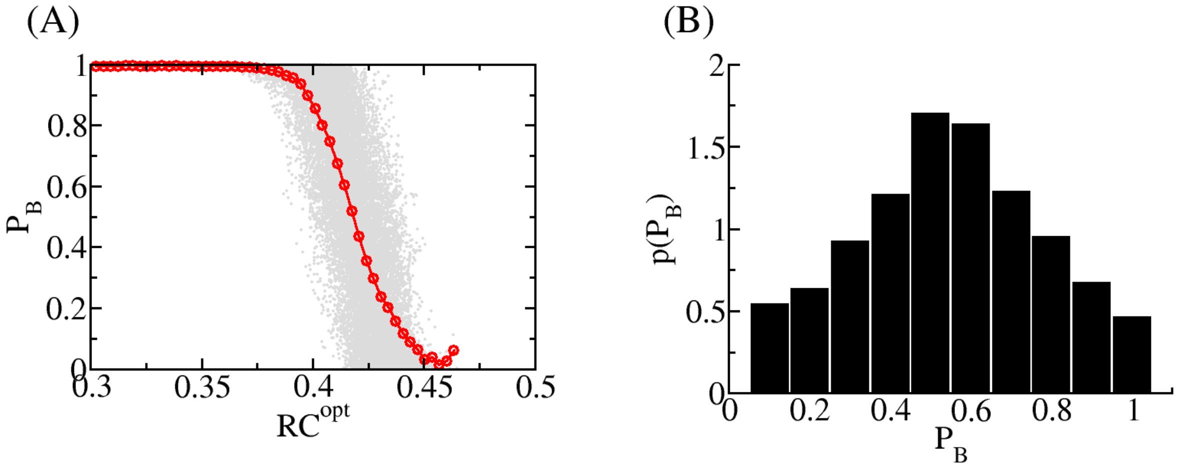


Figure S4: Committor analyses of the $O5 - CB10$ distance. (A) The averaged committor as a function of the $O5 - CB10$ distance (red line with circles). The committors of all configurations are shown as circles in grey. (B) Histogram of the committors for configurations with the $O5 - CB10$ distance between 0.417 and 0.4195.

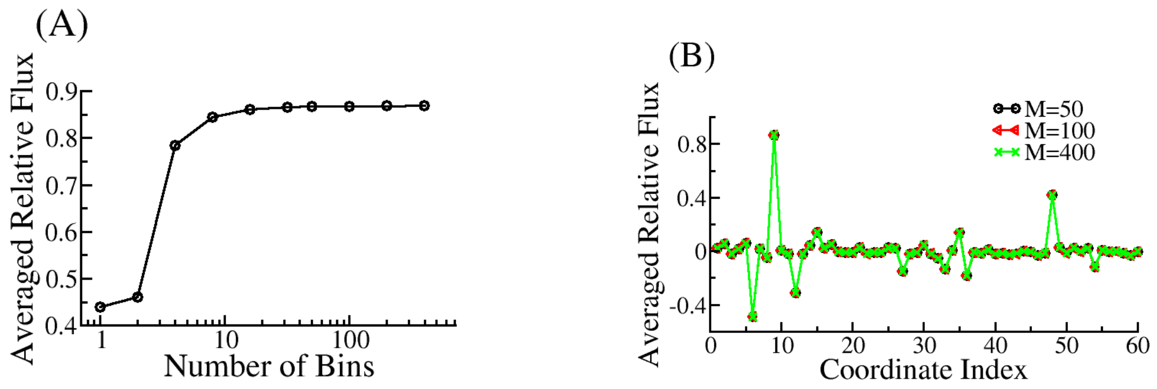


Figure S5: The averaged relative flux estimated with different number of bins. (A) The ARFs of $\phi$ as a function of $M$; (B) The ARFs of internal BAT coordinates calculated with M=50, 100, and 400.

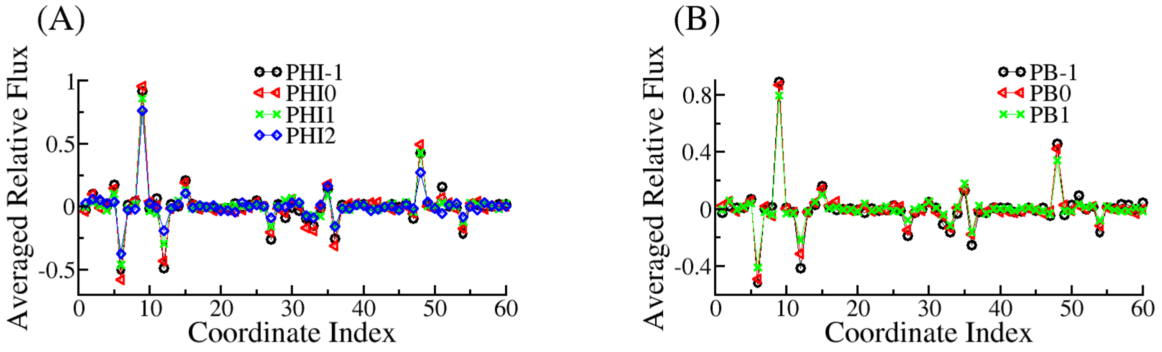


Figure S6: The averaged relative flux of internal BAT coordinates for different transition path sub-ensembles. (A) The ARFs for sub-ensembles defined by the committor; (B) The ARFs for sub-ensembles defined by $\phi$ and $\psi$.

Table S1: Information of all the internal BAT coordinates. Averaged relative flux (ARF), the rank by the magnitude of ARF (rank1), the coefficient in the optimized reaction coordinate (coef.), and the rank by the absolute value of the coefficient (rank2) for each coordinate are listed. “bon_A_B” is the bond between covalently bonded atoms $A$ and $B$; “ang_A_B_C” is the angle between a triplet of atoms “A-B-C”; “dih_A_B_C_D” is the dihedral angle between the “A-B-C” and “B-C-D” planes, where a “A-B-C” plane lies the “A-B” and “B-C” bonds. , Here, $A$, $B$, $C$ and $D$ are atom names. The coordinates $\phi$, $\psi$, and $\theta$ are dih_C4_N6_CA8_C14(9th), dih_N6_CA8_C14_N16(12th), and dih_O5_C4_N6_CA8(6th), respectively. See Fig. S1 for the index of each atom. The coefficients are rescaled and thus the coefficient of the coordinate with maximum magnitude is 1.

| index | name | ARF | rank1 | coef. | rank2 |
|---|---|---|---|---|---|
| 1 | bon_C4_N6 | 0.024 | 25 | -0.011 | 23 |
| 2 | ang_O5_C4_N6 | 0.057 | 12 | -0.017 | 17 |
| 3 | bon_C4_O5 | -0.019 | 35 | -0.012 | 20 |
| 4 | bon_N6_CA8 | 0.016 | 39 | -0.001 | 31 |
| 5 | ang_C4_N6_CA8 | 0.06 | 11 | -0.032 | 8 |
| 6 | dih_O5_C4_N6_CA8 | -0.49 | 2 | 0.477 | 2 |
| 7 | bon_CA8_C14 | 0.021 | 30 | 0.014 | 19 |
| 8 | ang_N6_CA8_C14 | -0.049 | 15 | 0 | 56 |
| 9 | dih_C4_N6_CA8_C14 | 0.869 | 1 | 1 | 1 |
| 10 | bon_C14_N16 | 0.004 | 54 | 0 | 60 |
| 11 | ang_CA8_C14_N16 | -0.022 | 27 | 0.018 | 15 |
| 12 | dih_N6_CA8_C14_N16 | -0.312 | 4 | 0.044 | 6 |
| 13 | bon_N16_CH18 | -0.021 | 31 | -0.005 | 29 |
| 14 | ang_C14_N16_CH18 | 0.045 | 17 | -0.022 | 12 |
| 15 | dih_CA8_C14_N16_CH18 | 0.14 | 7 | -0.062 | 5 |
| 16 | bon_CH18_H19 | 0.022 | 26 | 0.01 | 24 |
| 17 | ang_N16_CH18_H19 | 0.052 | 14 | 0.019 | 14 |
| 18 | dih_C14_N16_CH18_H19 | -0.004 | 55 | 0 | 35 |
| 19 | bon_CH18_H20 | -0.01 | 44 | -0.001 | 32 |
| 20 | ang_N16_CH18_H20 | -0.008 | 49 | 0 | 39 |
| 21 | dih_H19_N16_CH18_H20 | 0.026 | 22 | 0 | 51 |
| 22 | bon_CH18_H21 | -0.017 | 37 | 0.005 | 28 |
| 23 | ang_N16_CH18_H21 | -0.008 | 48 | 0 | 46 |
| 24 | dih_H19_N16_CH18_H21 | -0.007 | 51 | 0 | 37 |
| 25 | bon_N16_H17 | 0.025 | 23 | -0.012 | 21 |
| 26 | ang_C14_N16_H17 | 0.02 | 32 | 0 | 47 |
| 27 | dih_H17_N16_C14_CH18 | -0.149 | 6 | -0.027 | 9 |
| 28 | bon_C14_O15 | -0.019 | 34 | -0.006 | 25 |
| 29 | ang_CA8_C14_O15 | -0.01 | 46 | 0 | 53 |
| 30 | dih_O15_C14_CA8_N16 | 0.046 | 16 | -0.024 | 11 |

Continued on next page

Table S1: Information of all the internal BAT coordinates. Averaged relative flux (ARF), the rank by the magnitude of ARF (rank1), the coefficient in the optimized reaction coordinate (coef.), and the rank by the absolute value of the coefficient (rank2) for each coordinate are listed. "bon_A_B" is the bond between covalently bonded atoms $A$ and $B$; "ang_A_B_C" is the angle between a triplet of atoms "A-B-C"; "dih_A_B_C_D" is the dihedral angle between the "A-B-C" and "B-C-D" planes, where a "A-B-C" plane lies the "A-B" and "B-C" bonds. , Here, $A$, $B$, $C$ and $D$ are atom names. The coordinates $\phi$, $\psi$, and $\theta$ are dih_C4_N6_CA8_C14(9th), dih_N6_CA8_C14_N16(12th), and dih_O5_C4_N6_CA8(6th), respectively. See Fig. S1 for the index of each atom. The coefficients are rescaled and thus the coefficient of the coordinate with maximum magnitude is 1. (Continued)

| index | name | ARF | rank1 | coef. | rank2 |
|---|---|---|---|---|---|
| 31 | bon_CA8_H9 | -0.021 | 28 | -0.016 | 18 |
| 32 | ang_N6_CA8_H9 | -0.052 | 13 | 0 | 55 |
| 33 | dih_H9_CA8_N6_C14 | -0.133 | 9 | 0 | 54 |
| 34 | bon_CA8_CB10 | 0.007 | 50 | 0 | 58 |
| 35 | ang_N6_CA8_CB10 | 0.138 | 8 | -0.037 | 7 |
| 36 | dih_CB10_CA8_N6_C14 | -0.179 | 5 | 0.064 | 4 |
| 37 | bon_CB10_H11 | -0.006 | 52 | -0.019 | 13 |
| 38 | ang_CA8_CB10_H11 | -0.017 | 36 | 0 | 48 |
| 39 | dih_N6_CA8_CB10_H11 | 0.01 | 45 | 0 | 34 |
| 40 | bon_CB10_H12 | -0.019 | 33 | 0 | 59 |
| 41 | ang_CA8_CB10_H12 | -0.016 | 40 | 0 | 42 |
| 42 | dih_H11_CA8_CB10_H12 | -0.027 | 20 | 0 | 50 |
| 43 | bon_CB10_H13 | -0.017 | 38 | -0.001 | 33 |
| 44 | ang_CA8_CB10_H13 | 0.002 | 59 | 0 | 49 |
| 45 | dih_H11_CA8_CB10_H13 | -0.005 | 53 | 0 | 41 |
| 46 | bon_N6_H7 | -0.03 | 19 | -0.011 | 22 |
| 47 | ang_C4_N6_H7 | -0.013 | 42 | 0 | 43 |
| 48 | dih_H7_N6_C4_CA8 | 0.421 | 3 | -0.024 | 10 |
| 49 | bon_CH1_C4 | 0.027 | 21 | 0.006 | 26 |
| 50 | ang_CH1_C4_O5 | -0.013 | 41 | 0 | 52 |
| 51 | dih_CH1_C4_O5_N6 | 0.024 | 24 | -0.084 | 3 |
| 52 | bon_H0_CH1 | 0.001 | 60 | 0 | 57 |
| 53 | ang_H0_CH1_C4 | 0.021 | 29 | 0 | 44 |
| 54 | dih_H0_CH1_C4_O5 | -0.117 | 10 | 0 | 36 |
| 55 | bon_CH1_H2 | 0.008 | 47 | -0.001 | 30 |
| 56 | ang_H2_CH1_C4 | -0.002 | 58 | 0 | 45 |
| 57 | dih_H0_C4_CH1_H2 | -0.003 | 56 | 0 | 38 |
| 58 | bon_CH1_H3 | -0.011 | 43 | -0.006 | 27 |
| 59 | ang_H3_CH1_C4 | -0.033 | 18 | -0.017 | 16 |
| 60 | dih_H0_C4_CH1_H3 | -0.002 | 57 | 0 | 40 |

Table S2: Information of all the pairwise distances. "A_B" is the distance between atoms $A$ and $B$. The coefficients are rescaled and thus the coefficient of the coordinate with maximum magnitude is 1. See also Table S1 for the more explanation.

| index | name | ARF | rank1 | coef. | rank2 |
|---|---|---|---|---|---|
| 1 | CH1-C4 | 0.031 | 28 | 0.012 | 35 |
| 2 | CH1-O5 | 0 | 45 | 0 | 42 |
| 3 | CH1-N6 | -0.015 | 37 | -0.011 | 38 |
| 4 | CH1-CA8 | 0.067 | 19 | 0.043 | 14 |
| 5 | CH1-CB10 | -0.482 | 3 | -0.098 | 6 |
| 6 | CH1-C14 | 0.03 | 29 | 0.001 | 41 |
| 7 | CH1-O15 | -0.046 | 24 | 0.039 | 19 |
| 8 | CH1-N16 | 0.278 | 7 | -0.119 | 4 |
| 9 | CH1-C18 | 0.225 | 10 | 0 | 44 |
| 10 | C4-O5 | -0.023 | 33 | -0.022 | 30 |
| 11 | C4-N6 | 0.016 | 35 | -0.006 | 40 |
| 12 | C4-CA8 | 0.054 | 23 | -0.04 | 18 |
| 13 | C4-CB10 | -0.748 | 2 | -0.097 | 7 |
| 14 | C4-C14 | 0.058 | 22 | 0.144 | 3 |
| 15 | C4-O15 | 0.015 | 36 | -0.038 | 21 |
| 16 | C4-N16 | 0.172 | 12 | 0.062 | 13 |
| 17 | C4-C18 | 0.161 | 13 | 0.036 | 24 |
| 18 | O5-N6 | 0.042 | 25 | -0.03 | 27 |
| 19 | O5-CA8 | 0.064 | 20 | -0.184 | 2 |
| 20 | O5-CB10 | -0.815 | 1 | 1 | 1 |
| 21 | O5-C14 | 0.114 | 17 | -0.067 | 11 |
| 22 | O5-O15 | 0.143 | 14 | -0.033 | 25 |
| 23 | O5-N16 | -0.087 | 18 | -0.076 | 9 |
| 24 | O5-C18 | 0.011 | 41 | -0.036 | 23 |
| 25 | N6-CA8 | 0.026 | 30 | 0.042 | 16 |
| 26 | N6-CB10 | 0.123 | 16 | -0.112 | 5 |
| 27 | N6-C14 | -0.035 | 27 | -0.012 | 36 |
| 28 | N6-O15 | -0.213 | 11 | 0 | 45 |
| 29 | N6-N16 | 0.366 | 4 | 0.042 | 17 |
| 30 | N6-C18 | 0.289 | 5 | -0.042 | 15 |
| 31 | CA8-CB10 | 0.007 | 42 | -0.019 | 31 |
| 32 | CA8-C14 | 0.022 | 34 | 0.027 | 28 |
| 33 | CA8-O15 | -0.012 | 40 | -0.038 | 20 |
| 34 | CA8-N16 | 0.003 | 44 | 0.013 | 33 |
| 35 | CA8-C18 | 0.023 | 32 | 0.038 | 22 |
| 36 | CB10-C14 | 0.134 | 15 | -0.085 | 8 |
| 37 | CB10-O15 | 0.285 | 6 | 0.066 | 12 |
| 38 | CB10-N16 | -0.232 | 9 | 0 | 43 |
| 39 | CB10-C18 | -0.24 | 8 | -0.072 | 10 |
| 40 | C14-O15 | -0.024 | 31 | 0.033 | 26 |
| 41 | C14-N16 | 0.005 | 43 | -0.011 | 37 |
| 42 | C14-C18 | 0.04 | 26 | 0.027 | 29 |
| 43 | O15-N16 | 0.013 | 39 | -0.009 | 39 |
| 44 | O15-C18 | 0.063 | 21 | -0.016 | 32 |
| 45 | N16-C18 | -0.013 | 38 | -0.013 | 34 |
